# TurboID-based proximity labeling identifies novel germline proteins that maintain E granule integrity and small RNA homeostasis in *C. elegans*

**DOI:** 10.1101/2024.12.03.626736

**Authors:** Kun Li, Xuezhu Feng, Ke Wang, Xiaona Huang, Chengming Zhu, Xinya Huang, Quan Wen, Shouhong Guang, Xiangyang Chen

## Abstract

Germ granules are biomolecular condensates composed of RNA and proteins that play crucial roles in RNA metabolism and posttranscriptional gene regulation. *C. elegans* germ granules consist of at least seven distinct subcompartments, including P granules, Z granules, Mutator foci, SIMR foci, P-bodies, D granules and E granules. Among these condensates, the E granule, which is nonrandomly positioned within the germ granule, is required for the production of a specialized class of small interfering RNAs (siRNAs). However, the mechanisms underlying E granule formation and its functional significance remain largely unexplored. In this study, via the use of TurboID-based proximity labeling technology combined with an RNAi-based reverse genetic screen, we identified two novel components of the E granule, EGC-2/C27B7.5 and EGC-3/F59G1.8, which initiate E granule assembly. The depletion of EGC-2 or EGC- 3 disrupts the perinuclear localization of the EGO and PICS complexes, both of which are enriched in E granules and are required for E-class siRNA and piRNA biogenesis, respectively. Small RNAomic analyses revealed that both EGC-2 and EGC-3 promote the production of 5*’* E-class siRNA, whereas prohibiting piRNA maturation. Taken together, our results elucidate the roles of EGC-2 and EGC-3 in maintaining E granule integrity and small RNA homeostasis, and suggest that intracellular condensation may exert distinct regulatory effects on different biological processes. Additionally, the combination of proximity labeling technology and reverse genetic screening provides a robust strategy for studying the assembly of biomolecular condensates.

## Introduction

Biomolecular condensates are micron-scale membraneless organelles (MLOs) present in eukaryotic cells, common examples of which include processing bodies, stress granules, nucleoli, Cajal bodies and germ granules[1–3]. Current models posit that the formation of the biomolecular condensate is driven by liquid‒liquid phase separation (LLPS) via multivalent interactions between RNAs, intrinsically disordered proteins and RNA-binding proteins[1, 4]. These biomolecular condensates help cells spatiotemporally compartmentalize proteins and RNA molecules within distinct subcellular compartments to coordinate complicated RNA metabolism and gene expression processes[1, 5, 6]. Furthermore, many biomolecular condensates contain distinct immiscible subcompartments, forming multilayered architectures, which may regulate sequential RNA processing events; for example, distinct subdomains of the nucleolus participate in different ribosomal RNA processing steps[7–9]. However, the molecular mechanisms underlying the formation of these multilayered condensates and their cellular functions are still largely unknown.

Germ granules are RNA-rich biomolecular condensates that are anchored on the periphery of the nucleus and are thought to act as organizational hubs for posttranscriptional gene regulation[10–13]. Germ granules are widely present in many animals, including worms, flies, zebrafish, Xenopus and mice[5, 13–16]. In *C. elegans*, recent studies have shown that germ granules are divided into at least seven distinct subcompartments, including P granules, Z granules, Mutator foci, SIMR foci, D granules, E granules and P-bodies. The P granule is the first membraneless organelle found to be formed by liquid‒liquid phase separation and to exhibit liquid‒like behaviors, including fusion, dripping, and wetting[17]. P granules are thought to be major sites of mRNA regulation in germ cells and serve as hubs for self-/nonself RNA discrimination via siRNAs and the RNA interference (RNAi) machinery[13, 18]. PGL-1 is a germline-expressed protein that is a P granule component at all stages of development and is widely used as a marker protein of P granules[19–21]. Z granules, marked by the conserved SF1 helicase ZNFX-1, are reported to be adjacent to P granules, and regulate RNAi inheritance[21, 22]. Mutator foci (marked by MUT-16) and E granules (marked by ELLI-1) are two independent germ granule subcompartments that promote the production of siRNAs targeting distinct subsets of germline RNAs, respectively[23–26]. SIMR foci, marked by SIMR- 1, act downstream of the piRNA pathway to promote siRNA amplification via the Mutator complex and drive small RNA specificity for the nuclear Argonaute protein HRDE-1[27, 28]. P-bodies, which are marked by CGH-1 and are found to be located on top of P granules in pachytene cells, promote the segregation of germ granules into subcompartments and small RNA-based transgenerational gene silencing[29]. D granules, marked by DDX-19, are reported to concentrate between the zones of P granules and the nuclear pore complexes (NPCs), forming a tripartite architecture[30, 31]. Intriguingly, these immiscible germ granule subcompartments are not randomly ordered with respect to each other. For instance, many germ granules contain a single Z granule sandwiched between a P granule and a Mutator focus, forming ordered tri-condensate assemblages termed PZM[21]. SIMR foci are also found to be spatially organized, adjacent to Z granules and Mutator foci, and opposite P granules[27]. Moreover, a recent study discovered a toroidal P granule morphology in the mid- and late pachytene regions of the germline, which encircles the other germ granule compartments in a consistent exterior-to-interior spatial organization, further revealing the hierarchical organization of germ granules[32]. LOTUS domain proteins, including MIP-1, MIP-2 and LOTR-1, are reported to be required for the perinuclear assembly of particular germ granule compartments[22, 31, 33, 34]. Animals lacking MIP-1 and MIP-2 exhibit temperature-sensitive embryonic lethality, sterility, and mortal germlines, suggesting that the perinuclear anchoring of germ granules is essential for germ cell development[33, 34]. Interestingly, a recent study reported that a novel germline protein, HERD-1, regulates multiphase condensate immiscibility to mediate small RNA-driven transgenerational epigenetic inheritance, suggesting that the multiphasic architecture of the germ granule is crucial for gene expression regulation[35]. The current model posits that the multiphasic architecture of *C. elegans* germ granules helps cells organize numerous perinuclear proteins and coordinate highly sophisticated perinuclear gene regulation networks, especially small RNA-based gene regulatory pathways[13, 18, 36].

Although the molecular mechanisms of the assembly of biomolecular condensates have been widely investigated, the cellular functions of many biomolecular condensates and the biological significance of the positioning of proteins in these condensates are still largely unknown. Deciphering the genetic requirements of the assembly of particular condensates may help elucidate the biological functions of these membraneless organelles. For example, RNA-binding proteins such as GLH-1, PGL-1/PGL-3, and MEG-3/MEG-4, play important roles in the formation of P granules[37–42]. The depletion of these assembly factors usually leads to disordered expression of RNAomes, including small RNAs and mRNAs[43–47], suggesting that P granules are crucial for the maintenance of RNA homeostasis. The intrinsically disordered protein MUT-16 functions as a scaffold and promotes phase separation of Mutator foci, the depletion of which blocks the production of Mutator class siRNAs and results in defects in fertility[23, 24]. Another intrinsically disordered protein, PID-2/ZSP-1, which was reported to be needed for Z granule homeostasis, is required for heritable piRNA-induced silencing and germline immortality[48, 49]. Our recent works identified the E granule as a novel subcompartment of *C. elegans* germ granules and revealed that both the PICS/PETISCO complex (PICS-1/TOFU-6/PID-1/ERH-2) and the EGO complex (ELLI-1/EGC-1/EGO- 1/DRH-3/EKL-1) are enclosed in the E granule[26, 31, 50]. The PICS complex is required for piRNA processing to produce mature piRNAs[50, 51] and the EGO complex is required for the generation of a specialized subset of siRNAs[26, 52]. Yet, the cellular function of the perinuclear positioning of these two complexes is largely unknown. Identification of proteins that mediate E granule assembly and deciphering the E granule assembly process may help reveal how E granules coordinate multiple biological processes and why cells enclose different complexes in the same germ granule compartment.

In this study, we utilized TurboID enzyme-catalyzed proximity labeling technology to identify additional components of the E granule in *C. elegans* germline. This approach led to the identification of 139 candidate proteins potentially localized to E granules. Through an RNAi-based reverse genetic screen, we successfully identified two uncharted proteins, C27B7.5/EGC-2 (<u>E</u> <u>g</u>ranule <u>c</u>omponent-2) and F59G1.8/EGC-3 (<u>E</u> <u>g</u>ranule <u>c</u>omponent-3), which facilitate the perinuclear localization of both the PICS and the EGO complexes. We demonstrated that these proteins are enclosed in the E granule and exclusively promote E granule assembly. We further performed small RNAomic analyses and revealed that both EGC- 2 and EGC-3 promote the production of 5’ E-class siRNA, whereas limiting piRNA biogenesis.

Additionally, we found that EGC-2 and EGC-3 contribute to the RNA interference (RNAi) response. Taken together, our results identify key factors involved in E granule assembly that are essential for maintaining small RNA homeostasis, and suggest that intracellular condensation may exert distinct regulatory effects on different biological processes.

## Results

### The perinuclear accumulation of the EGO complex and PICS complex is independent of each other

The current model posits that the *C. elegans* germ granule is composed of multiple subcompartments (Figure. 1A) [13, 31]. In previous studies, we identified a novel germ granule compartment, the E granule, which encloses the EGO complex (consisting of ELLI-1, EGC-1, EGO-1, EKl-1 and DRH-3) and the PICS complex (consisting of PICS-1, TOFU-6, PID-1 and ERH-2) (Figure. 1B) [26, 31, 50, 53].

**Figure 1.**
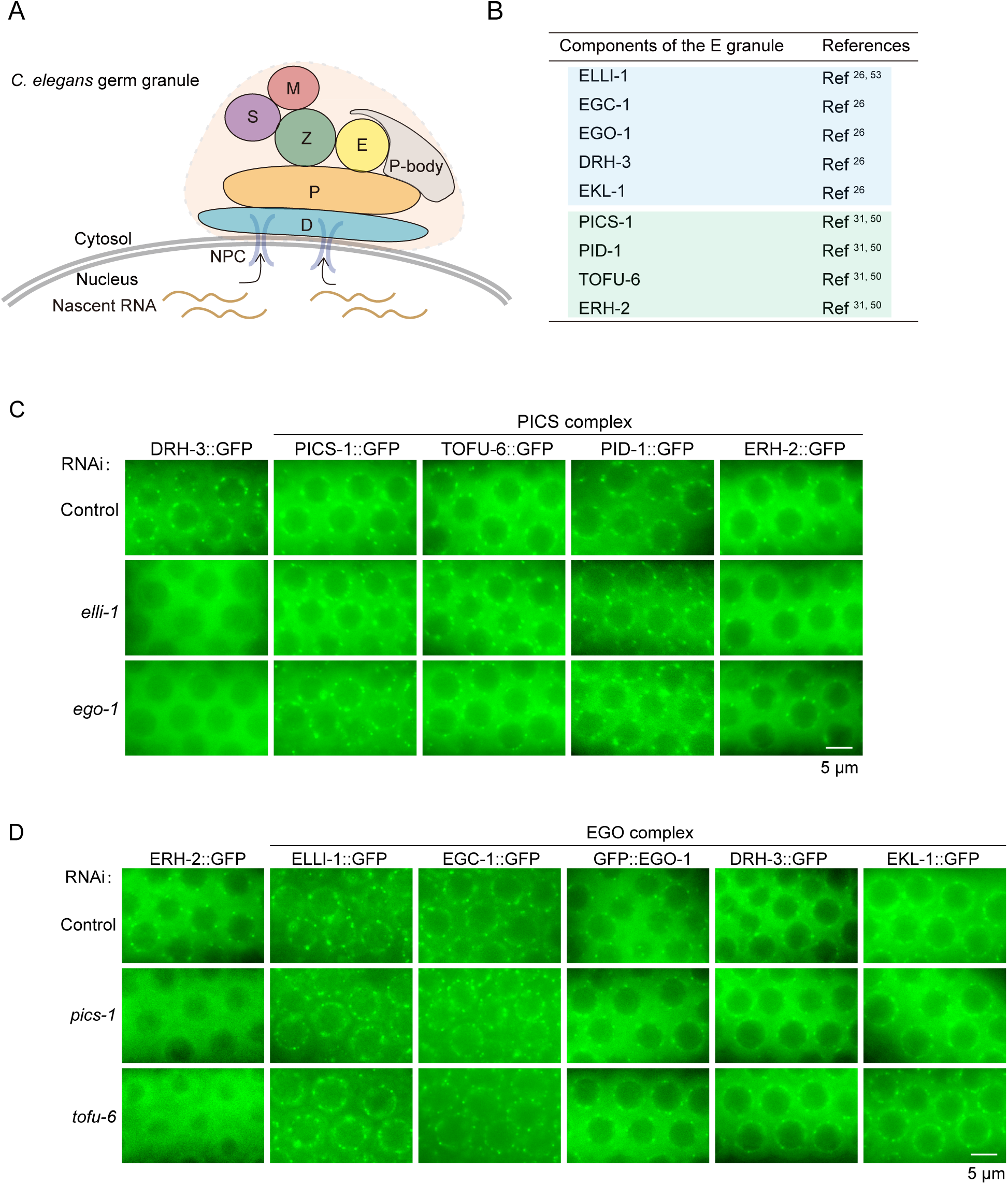
The perinuclear accumulation of the EGO complex and PICS complex is independent of each other. (A) Schematic representation of the germ granule architecture in *C. elegans*. (B) Summary of nine identified E granule components and corresponding references. (C) Fluorescence microscopy of the GFP-tagged PICS complex components in adult germ cells following RNAi targeting *elli-1* and *ego-1*. RNAi knockdown of *elli-1* and *ego-1* disrupted DRH-3::GFP foci, whereas PICS complex localization remained unaffected. (D) Fluorescence micrographs of the GFP-tagged EGO complex components in adult germ cells following RNAi targeting *tofu-6* and *pics-1*. RNAi knockdown of *tofu-6* and *pics-1* affected ERH-2::GFP foci but did not disturb the perinuclear localization of the EGO complex. All images are representative of more than three animals.

We previously found that ELLI-1 and EGC-1 are essential for perinuclear accumulation of the EGO module (comprising EGO-1, EKL-1 and DRH-3). We further investigated which components of the EGO complex may play the most upstream role in their perinuclear accumulation by examining whether the EGO module factors promote the perinuclear accumulation of ELLI-1 or EGC-1. Feeding RNAi targeting *ego-1* likely reduced the expression level of ELLI-1::GFP, yet a considerable number of ELLI-1 foci can still be observed in germ cells; RNAi knockdown of *ego-1* completely disrupted perinuclear EGC-1 foci, suggesting that EGO-1 is required for the perinuclear accumulation of EGC-1 (Figure. S1A); the DRH-3 foci and EKL-1 foci were also significantly disrupted upon *ego-1* knockdown, and the residual foci of DRH-3 and EKL-1 colocalized with tagRFP::MUT-16 (Figure. S1B, S1C). The knockdown of DRH-3 via an auxin-inducible degron system also elicited the diffusion of both EGC-1 and EGO-1 in germ cells, yet did not affect the perinuclear enrichment of ELLI-1 (Figure. S1A). Together, these results suggest that ELLI-1 plays the most upstream role in the assembly of the EGO complex in E granules.

The PICS/PETISCO complex, which stabilizes the PUCH complex and facilitates 5’ trimming of piRNA precursors, was enriched in the E granule[31]. We examined whether the EGO complex components, ELLI-1 and EGO-1, are required for the E granule accumulation of the PICS complex as well. Strikingly, RNAi knockdown of *elli-1* and *ego-1* did not disrupt the formation of the PICS foci (Figure. 1C). The PICS foci disassembled in the absence of *pics- 1* or *tofu-6* [50]. RNAi knockdown of *pics-1* and *tofu-6* disrupted the perinuclear foci of ERH- 2, yet, did not affect the formation of the EGO complex foci (Figure. 1D).

Taken together, the results suggested that the perinuclear accumulation of the EGO and PICS complexes was likely independent of each other, hinting the presence of certain master organizers that may act upstream to assemble E granules.

### Identification of E granule components via TurboID-based proximity labeling

To identify additional proteins that promote E granule assembly, we sought to investigate the protein components of E granules. TurboID is an engineered biotin ligase that uses ATP to convert biotin into biotin-AMP, a reactive intermediate that covalently labels proximal proteins in living cells [34, 54, 55]. We employed this biotin ligase-based proximity labeling method to label E granule proteins. We used CRISPR/Cas9 to introduce the TurboID sequence into the genomic loci of *elli-1* (Figure. 2A, left panel). The insertion of the TurboID sequence was confirmed by genotyping via PCR and DNA sequencing (Figure. 2A, right panel).

**Figure 2.**
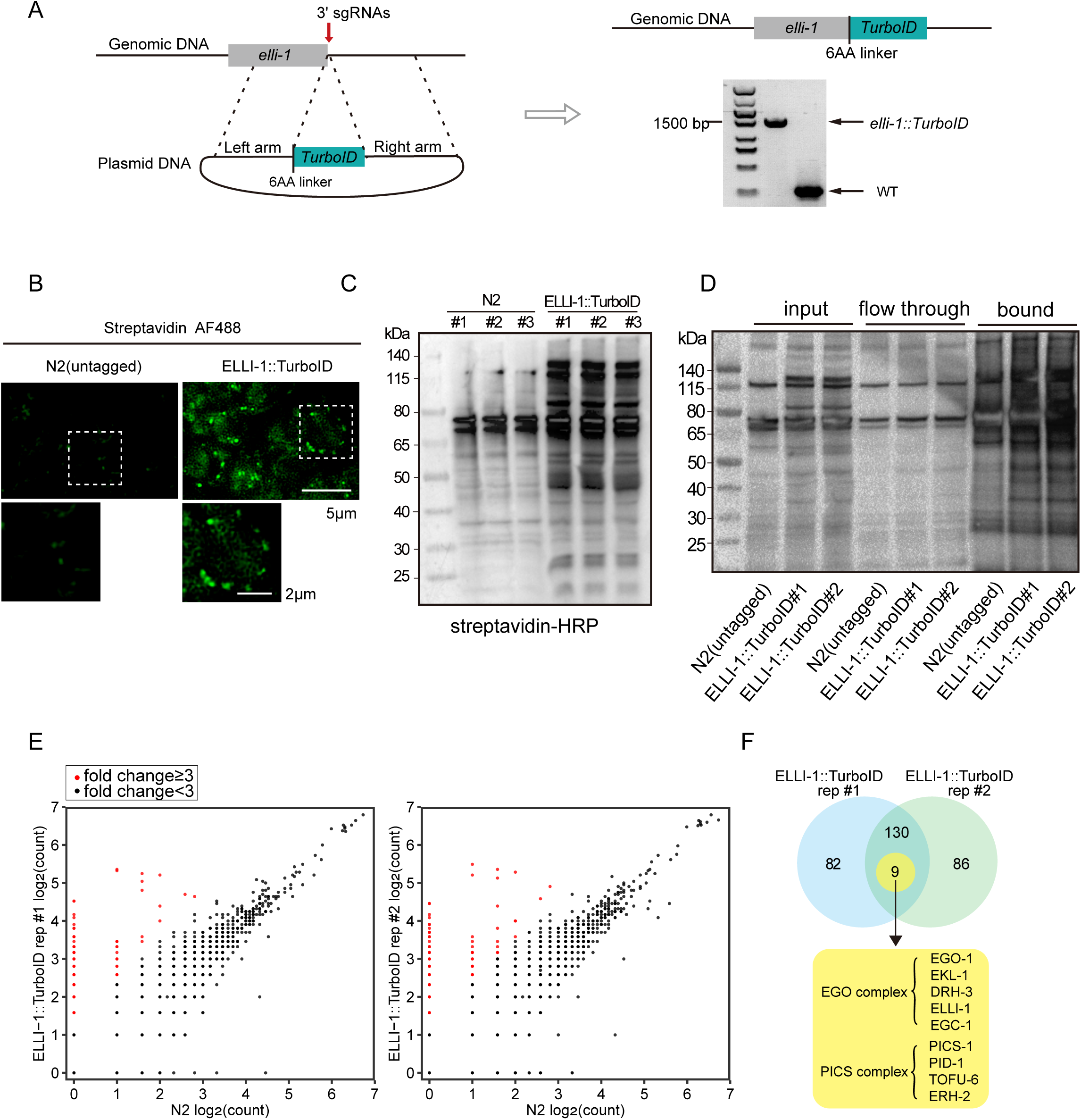
Identification of E granule proteins via TurboID-based proximity labeling. (A) Schematic of the construction of the *elli-1::TurboID* proximity labeling transgene. The TurboID coding sequence, fused with a 6-amino acid linker, was inserted upstream of the stop codon in the *elli-1* genomic locus via CRISPR/Cas9 technology. The correct insertion was confirmed through genotyping and sequencing. (B) Fluorescence micrographs of streptavidin-Alexa Fluor 488-stained germ cells from untagged N2 and *elli-1*::TurboID animals. (C) Western blot image showing streptavidin-horseradish peroxidase (HRP) labeling of whole-animal lysates from wild-type and *elli-1::TurboID* animals across three independent biological replicates. (D) Streptavidin affinity pull-down efficiency shown by streptavidin-HRP blotting. Biotinylated proteins were extracted from the input via a specialized buffer (see methods), showing enrichment in the pull-down fraction, with minimal endogenously biotinylated proteins detected in the flow-through. The enrichment of biotinylated proteins was confirmed in one replicate for the untagged strains and two replicates for the *elli-1::TurboID* strains. (E) Scatter plots showing the fold changes in unique peptide counts derived from mass spectrometry data, comparing *elli-1::TurboID* to wild-type animals, with a pseudocount of 1 to accommodate zeros. Proteins with a fold change ≥3 (indicated by red dots) were artificially classified as putative E granule components, whereas those with a fold change <3 are shown as black dots. (F) Venn diagram showing 139 proteins identified in both TurboID-based proximity labeling replicates, including components of the EGO and PICS complexes.

We first assessed if the tagged *elli-1* allele generates functional proteins by examining the feeding RNAi response of animals expressing ELLI-1::TurboID proteins. Loss of ELLI-1 results in defects in feeding RNAi responses targeting germline genes[26]. Compared with wild-type animals, animals expressing ELLI-1::TurboID proteins responded normally to feeding RNAi targeting *pos-1*, *mex-3* and *mex-5* (Figure. S2A). To determine whether E granules are normally assembled in the *elli-1::TurboID* strain, we examined the subcellular localization of EGC-1, a well-characterized E granule protein. Like in wild-type animals, EGC- 1::GFP primarily accumulated in perinuclear foci in *elli-1::TurboID* animals (Figure. S2B). Together, the results suggest that the modified *elli-1* gene encodes functional proteins.

We then assayed the biotinylation of proteins by immunofluorescence staining and streptavidin blotting to assess the enzyme activity of TurboID. First, we stained dissected gonads with fluorescently labeled streptavidin to examine the localization of biotinylated proteins in germ cells. Cytoplasmic signals in the stained wild-type gonads were barely detectable, whereas pronounced perinuclear signals of biotinylated proteins were observed in *elli-1::TurboID* gonads (Figure. 2B). Furthermore, streptavidin–horseradish peroxidase blot analysis of lysed adult animals revealed a dramatically increased presence of biotinylated proteins in *elli-1::TurboID* samples compared with untagged control samples (Figure. 2C). These results suggested that ELLI-1::TurboID can be applied to label proteins in living germ cells.

We next conducted streptavidin affinity pull-down assays to enrich TurboID-biotinylated proteins. Blotting results confirmed that biotinylated proteins were efficiently enriched in the pull-down samples, with only minimal endogenously biotinylated proteins remaining in the flow-through (see methods) (Figure. 2D). Biotinylated proteins from the untagged control and *elli-1::TurboID* strains were enriched in one and two independent biological replicates, respectively, and were identified via mass spectrometry. Mass spectrometry analysis revealed that 139 proteins were significantly enriched in the ELLI-1::TurboID samples compared with the untagged control (fold change of unique peptide counts ≥ 3) (Figure. 2E), as listed in Table S1. The candidate list included all known components of the EGO complex (ELLI-1, EGC-1, EGO-1, DRH-3 and EKL-1) and the PICS complex (TOFU-6, PID-1, PICS-1 and ERH-2), suggesting that TurboID proximity labeling is effective for revealing proteins within E granules (Figure. 2F).

### Candidate-based RNAi screening identifies two proteins, EGC-2 and EGC-3, that promote the formation of perinuclear EGO foci and PICS foci

To identify uncharted E granule components involved in E granule assembly, we then performed RNAi-based genetic screening to pinpoint proteins that facilitate the formation of perinuclear ELLI-1 foci. From the 139 candidate proteins, we first refined the list by excluding proteins that were reported to be not enriched in the germline or localized to particular germ granule compartments other than the E granule. We subsequently knocked down 40 genes from the narrowed list in *elli-1::gfp* animals by feeding them bacteria expressing related dsRNAs to determine whether the depletion of these genes disrupted ELLI-1 foci formation (Figure. 3A). Two genes, *c27b7.5* and *f59b1.8*, were identified as essential for ELLI-1 foci formation (Figure. 3B). based on the data described above, *c27b7.5* and *f59b1.8* were named <u>E</u> <u>g</u>ranule <u>c</u>omponent-2 (*egc-2*) and <u>E</u> <u>g</u>ranule <u>c</u>omponent-3 (*egc-3)*, respectively.

**Figure 3.**
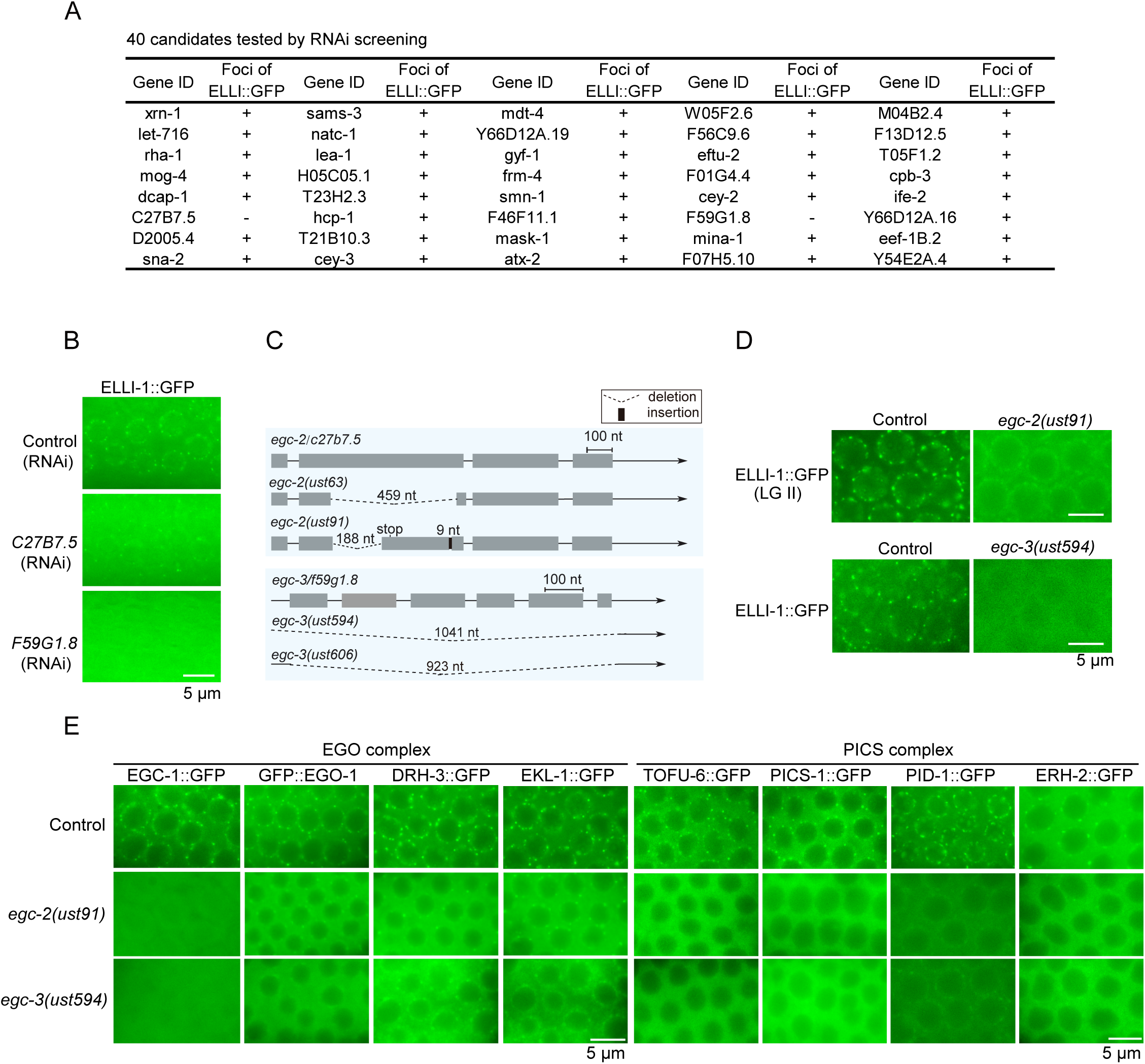
Candidate-based RNAi screening identifies two proteins, EGC-2 and EGC-3, that promote the formation of perinuclear EGO foci and PICS foci. (A) A summary table listing 40 candidate genes evaluated via RNAi-based reverse genetic screening. The impact on ELLI-1::GFP foci was assessed by feeding RNAi targeting these candidates; “-” denotes disruption of ELLI-1::GFP foci, whereas “+” indicates no effect. (B) Fluorescence micrographs of ELLI-1::GFP in pachytene germ cells upon RNAi targeting *C27B7.5* and *F59G1.8*. (C) Sequences of *egc-2/c27b7.5* and *egc-3/f59g1.8* alleles generated via a multiple sgRNA-guided CRISPR/Cas9 system. Deletions and insertions are drawn to scale. Premature termination codons caused by frameshift mutations are indicated. (D) Pachytene germ cells from dissected gonads expressing ectopic ELLI-1::GFP (LG II) or *in situ* ELLI-1::GFP (LG Ⅳ) in the indicated animals. (E) Fluorescence micrographs of the GFP-tagged components of the EGO and PICS complexes in the adult germlines of *egc-2(-)* and *egc-3(-)* animals. The loss of EGC- 2 or EGC-3 resulted in the disruption of perinuclear localization of both the EGO and the PICS complexes. All images are representative of more than three animals.

To further confirm that EGC-2/C27B7.5 and EGC-3/F59G1.8 promote ELLI-1 foci assembly, we generated mutant alleles of *egc-2* and *egc-3* via multiple sgRNAs guided CRISPR/Cas9 technology (Figure. 3C)[56]. Using genetic crosses, we generated *egc-2(-)* and *egc-3(-)* animals expressing ELLI-1::GFP. Consistent with the RNAi results, ELLI-1 foci were significantly disrupted in these mutant animals (Figure. 3D). Together, these results suggest that both EGC-2 and EGC-3 are required for ELLI-1 foci formation in germ cells.

Next, we examined whether EGC-2 and EGC-3 are required for the perinuclear localization of other components of the EGO and PICS complexes. In both *egc-2* and *egc-3* mutants, the perinuclear foci of EGC-1, TOFU-6, PICS-1 and ERH-2 were all abolished (Figure. 3E). The perinuclear foci of PID-1 were also substantially disrupted in *egc-2* and *egc-3* mutants, although few PID-1 aggregates remained observable (Figure. 3E). The localization of EGO module factors (EGO-1, DRH-3 and EKL-1) to perinuclear foci was significantly reduced in both *egc-2(-)* and *egc-3(-)* animals; however, residual EGO module-marked foci were still noticeable (Figure. 3E). These results suggest that EGC-2 and EGC-3 are required for perinuclear localization of both the EGO and PICS complexes.

### EGC-2 and EGC-3 are enriched in the E granule and promote E granule assembly

The assembly of many biomolecular condensates is driven by intrinsically disordered/low-complexity proteins[1]. Both EGC-2 and EGC-3 possess low-complexity domains, hinting that these proteins might facilitate E granule assembly (Figure. 4A, 4B). To investigate the expression patterns and subcellular localization of EGC-2 and EGC-3, we constructed a single-copy *gfp::3xflag*-tagged *egc-2* transgene using the MosSCI system[57] and introduced a *gfp::3xflag* epitope into the endogenous *egc-3* gene via CRISPR/Cas9 technology. Both proteins were expressed throughout the germline and predominantly accumulated in perinuclear foci in germ cells, similar to other E granule proteins (Figure. S3A)[26]. Yet, unlike previously reported E granule proteins[26], EGC-2 and EGC-3 were expressed and enriched mainly in cytoplasmic condensates in early embryos, implying that EGC-2 and EGC-3 may exercise cellular functions independent of the PICS and EGO complexes during embryogenesis (Figure. S3B).

**Figure 4.**
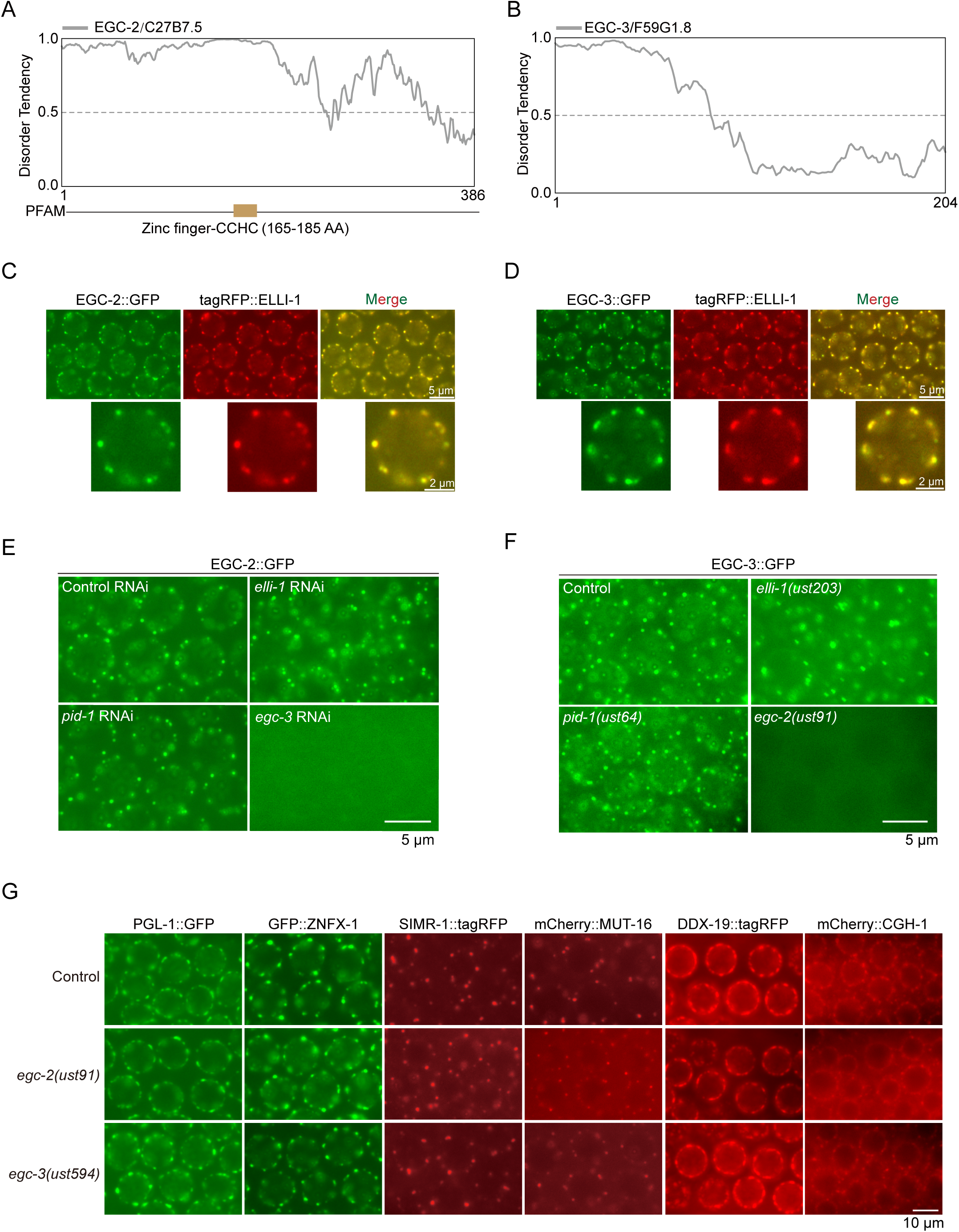
EGC-2 and EGC-3 are enriched in the E granule and promote E granule assembly. (A and B) Graphs showing the disordered tendency of EGC-2/C27B7.5 and EGC- 3/F59G1.8 as analyzed by the AIUPred program[89]. AIUPred provides a scoring system that assesses the disordered tendency of each amino acid position along the sequence, with scores ranging from 0 to 1. Residues with scores exceeding 0.5 are classified as disordered, whereas those below this threshold are classified as ordered. (C and D) Fluorescence micrographs of pachytene germ cells expressing the indicated fluorescent proteins. Both EGC-2::GFP and EGC-3::GFP colocalized with tagRFP::ELLI-1. (E and F) Fluorescence micrographs of EGC- 2::GFP or EGC-3::GFP in pachytene germ cells from the indicated animals. Depletion of ELLI- 1 or PID-1 did not disturb the perinuclear localization of EGC-2 and EGC-3. However, depletion of EGC-2 and EGC-3 mutually disrupted each other’s perinuclear localization. (G) Fluorescence micrographs of adult germ cells expressing PGL-1::GFP (marking P granule), GFP::ZNFX-1 (marking Z granule), SIMR-1::tagRFP (marking SIMR foci), mCherry::MUT-16 (marking Mutator foci), DDX-19::tagRFP (marking D granule) or mCherry::CGH-1 (marking P body) in the indicated animals. Depletion of either EGC-2 or EGC-3 did not disturb the perinuclear localization of the other six germ granule subcompartments. All images are representative of more than three animals.

To assess whether EGC-2 and EGC-3 localize to the E granule, we used genetic crosses to generate animals expressing combinations of tagRFP::ELLI-1 and EGC-2::GFP or EGC- 3::GFP. We found that both EGC-2::GFP and EGC-3::GFP colocalized with tagRFP::ELLI-1, suggesting that these two proteins are E granule components (Figure. 4C, 4D).

We further examined whether the EGO and PICS complexes regulate the formation of perinuclear EGC-2::GFP and EGC-3::GFP foci. Knockdown of *elli-1* or *pid-1* did not disrupt the perinuclear localization of EGC-2 or EGC-3, indicating that EGC-2 and EGC-3 function upstream of other E granule components in E granule assembly (Figure. 4E, 4F). Additionally, the perinuclear localization of EGC-2 and EGC-3 was disrupted in each other’s mutants, indicating that their perinuclear localization is mutually dependent (Figure. 4E, 4F).

The current model posits that *C. elegans* germ granules in pachytene cells consist of at least seven subcompartments. Thus, we examined whether EGC-2 and EGC-3 participate in the assembly of other germ granule compartments. Genetic crosses were performed to generate *egc-2* and *egc-3* mutant animals expressing fluorescent markers for each subcompartment. The results showed that the mutations specifically affected E granule assembly, without disrupting the formation of P granules (marked by PGL-1::GFP), Z granules (marked by GFP::ZNFX-1), SIMR foci (marked by SIMR-1::tagRFP), Mutator foci (marked by mCherry::MUT-16), D granules (marked by DDX-19::tagRFP) or P-bodies (marked by mCherry::CGH-1) (Figure. 4G).

Overall, these results reveal that EGC-2 and EGC-3 are dedicated E granule components that exclusively promote E granule assembly, with no detectable impact on the formation of other germ granule subcompartments.

### EGC-2 and EGC-3 prohibit piRNA production

The PICS/PETISCO complex engages in piRNA processing, the depletion of which blocks the production of mature piRNAs[50, 51]. Since the components of the PICS complex are all enriched in the E granule and their perinuclear localization requires both EGC-2 and EGC-3, we then examined whether the localization of the PICS complex in the germ granule is essential for piRNA processing. We extracted total RNA from wild-type, *egc-2(-)* and *egc-3(-)* animals, performed 5*’* phosphate-dependent small RNA sequencing and quantified each piRNA. The 5*’* phosphate-dependent small RNA sequencing method excludes secondary siRNA signals that have 5*’* triphosphate ends, and thus boosts the relative piRNA signals in the sequencing results. Surprisingly, the depletion of EGC-2 or EGC-3 did not inhibit the accumulation of mature piRNAs (Figure. 5A). In contrast, piRNA expression levels were modestly elevated in *egc-2(-)* and *egc-3(-)* animals, suggesting a potential role for EGC-2 and EGC-3 in limiting piRNA production (Figure. 5A).

**Figure 5.**
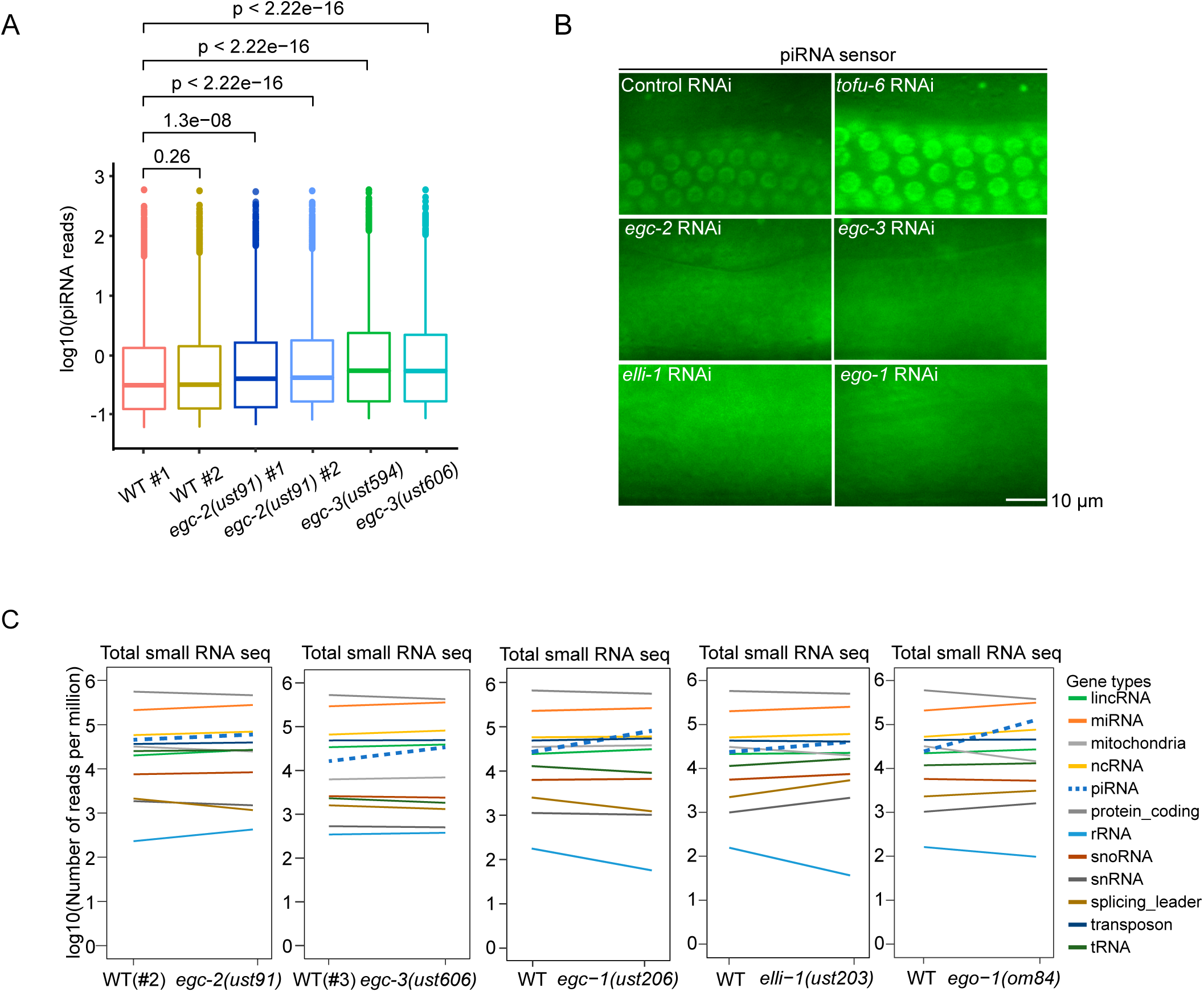
EGC-2 and EGC-3 prohibit piRNA production. (A) Boxplot showing normalized piRNA reads in *egc-2(-)*, *egc-3(-)* and wild-type animals at the young adult stage. Statistical significance was assessed via two-tailed *t* test, with the results indicated above the boxplot. The median piRNA abundance is shown as a solid black line. piRNA expression was significantly upregulated in *egc-3(-)* animals but slightly increased in *egc-2(-)* animals. (B) Fluorescence micrographs of piRNA sensor expression in the indicated RNAi-treated animals. Depletion of tofu-6 via RNAi served as a positive control, resulting in the desilencing of the piRNA sensor in the germline. RNAi knockdown of *egc-2*, *egc-3*, *elli-1* and *ego-1* further silenced the piRNA sensor. (C) Deep sequencing analysis of total small RNAs in the indicated animals. The blue dashed line indicates piRNAs.

To further confirm the role of EGC-2 and EGC-3 in piRNA production, we evaluated the activity of the piRNA-mediated germline surveillance pathway in *egc-2* and *egc-3* mutants via a piRNA sensor assay. The current model posits that piRNAs mediate the genome-wide surveillance of germline transcripts to silence non-self RNA transcripts in *C. elegans*[58–61]. In this assay, the GFP-labeled piRNA sensor transgene is typically silenced and only reveals a weak fluorescent signal in wild-type germlines but becomes desilenced when piRNA production is impaired or when piRNA-mediated gene silencing is disrupted[51, 62, 63]. TOFU-6 is required for piRNA production[50, 51]. RNAi-mediated depletion of TOFU-6 resulted in desilencing of the piRNA sensor (Figure. 5B). Yet, knockdown of *egc-2* or *egc-3* in the piRNA sensor strains further enhanced the silencing of the sensor, further supporting that EGC-2 or EGC-3 limits the production of piRNAs (Figure. 5B).

Previous studies identified EGO-1 and EKL-1 as suppressors of piRNA expression[64]. Since EGC-2, EGC-3, EGO-1 and EKL-1 are all components of the E granule, we investigated whether defects in E granule assembly or the perinuclear localization of the EGO complex affect piRNA levels. We sequenced the total small RNAs in *egc-2(-)* and *egc-3(-)* animals via a 5*’* phosphate-independent method and reanalyzed our published sequencing datasets[26]. Intriguingly, we found that the depletion of E granule proteins, including EGC-2, EGC-3, EGC- 1, ELLI-1 and EGO-1, other than the PICS complex, led to a general upregulation of piRNA levels (Figure. 5C). Consistently, knockdown of *elli-1* or *ego-1* by RNAi in germ cells resulted in further silencing of piRNA sensors (Figure. 5B).

Together, these results suggest that both EGC-2 and EGC-3 limit piRNA biogenesis. The perinuclear localization of the PICS complex in the E granule appears to play a role in regulating the activity of the piRNA-based gene silencing pathway, which might prevent excessive piRNA silencing.

### EGC-2 and EGC-3 promote the production of 5*’* E-class siRNA

Previous studies have shown that EGC-1 and ELLI-1, which facilitate perinuclear accumulation of the EGO module, are essential for 5*’* E-class siRNA production[26]. We investigated whether EGC-2 and EGC-3 also play a role in 5*’* E-class siRNA generation. We analyzed total small RNAs from wild-type, *egc-2(-)* and *egc-3(-)* animals. 22G RNAs were mapped to the *C. elegans* genome, and the number of siRNAs complementary to each *C. elegans* germline gene was quantified. E-class siRNAs were generally reduced in both *egc-2* and *egc-3* mutants, although the reduction of siRNAs in *egc-2* mutants was less pronounced than those in *egc-3* mutants (Figure. 6A, 6B).

**Figure 6.**
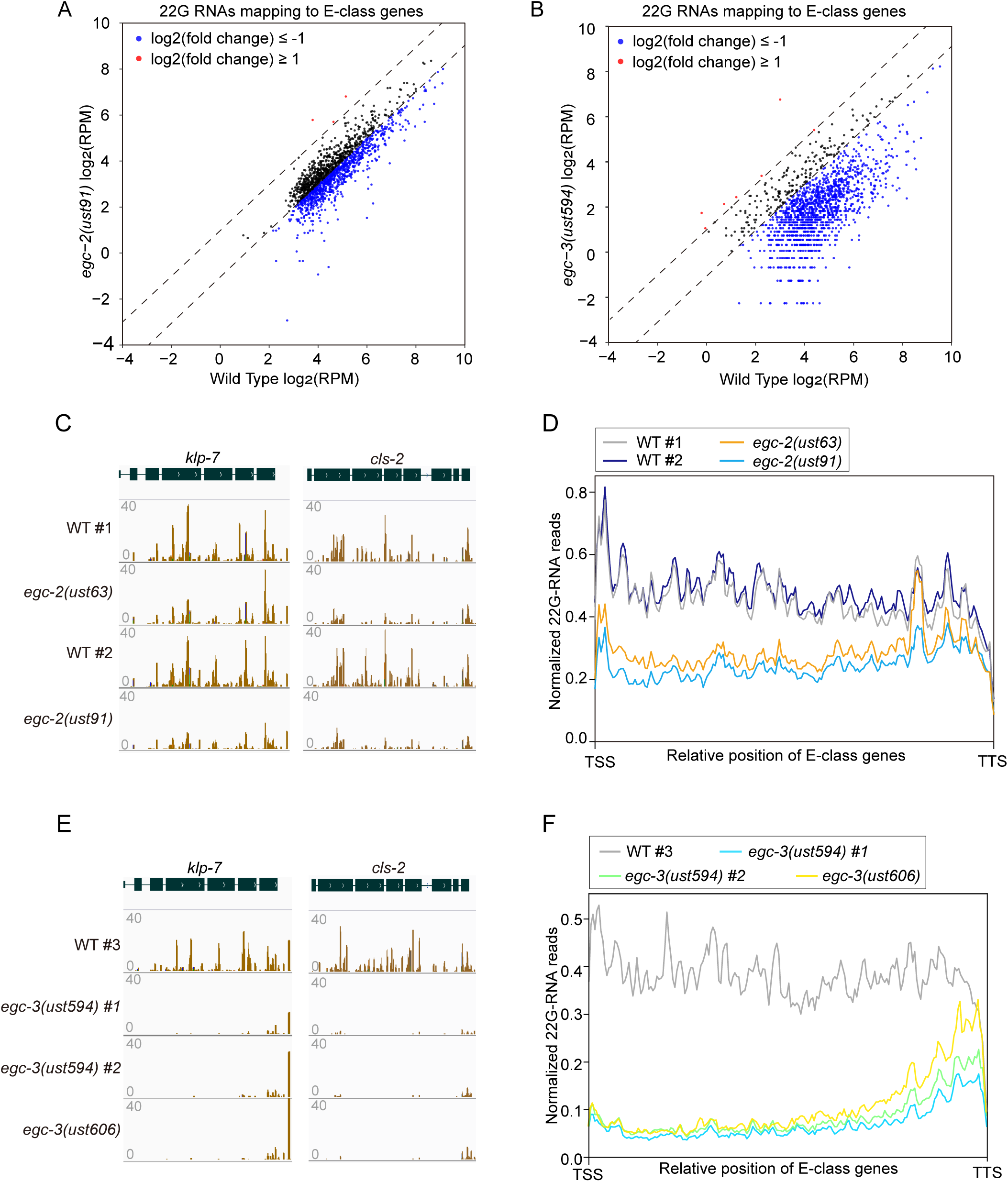
EGC-2 and EGC-3 promote the production of 5*’* E-class siRNA. (A and B) Scatter plots showing gene-by-gene comparisons of normalized siRNA abundances. siRNAs from wild-type, *egc-2(-)* and *egc-3(-)* animals were sequenced via a 5*’* phosphate-independent method. 22G RNAs were mapped to the *C. elegans* genome, and the number of reads complementary to each *C. elegans* gene was quantified. A cutoff criterion of a 2-fold change was applied to identify differentially expressed siRNAs. Genes with upregulated and downregulated siRNAs are shown in red and blue, respectively. (C and E) Normalized 22G RNA read distribution across E-class siRNAs targeting *klp-7* and *cls-2* in the indicated animals. Additional examples of E-class genes can be found in Figure S5. (D and F) Metaprofile analysis showing the distribution of normalized 22G RNA reads (RPMs) along E-class genes in the indicated animals.

The E-class siRNAs reduced in *egc-2* mutants were predominantly mapped to the 5*’* portion of the E-class genes. For instance, in wild-type animals, siRNAs targeting *klp-7, cls-2*, *F01G4.4*, *tebp-2, csr-1* and *hcp-1* were evenly distributed across the length of their target mRNAs. However, in *egc-2* mutants, siRNAs mapping to the 5*’* portions were generally reduced by more than half, whereas 3*’* end siRNAs remained relatively unaffected (Figure. 6C, S4A). Metagene analysis of 22G RNAs along E-class genes showed that the siRNAs mapping to the 5*’* portion, but not the 3*’* end siRNAs, of E-class genes was decreased in *egc-2* mutants (Figure. 6D). EGC-2 was not required for the production of M-class siRNAs, which are dependent on the Mutator protein, MUT-16 (Figure. S4B, S4C).

In *egc-3* mutants, the reduction of E-class siRNAs mapping to E-class genes was even more significant than in *egc-2* mutants, resembling the effects observed in *egc-1* and *elli-1* mutants[26]. For example, siRNAs mapping to the 5*’*, but not 3*’*-most, portions of *klp-7, cls-2*, *F01G4.4*, *tebp-2, csr-1* and *hcp-1* were almost entirely absent in *egc-3* mutants (Figure. 6E,S4D). Metagene analysis showed a substantial loss of siRNAs mapping to the 5’ portions of E-class genes in *egc-3(-)* animals, whereas 3’ end siRNAs also remained unaffected (Figure. 6F). Like EGC-2, EGC-3 was not involved in M-class siRNA production (Figure. S4E, S4F). These results suggest that while EGC-3 plays a role in E granule assembly, it might also participate in the amplification of siRNAs along E-class mRNAs (see discussion).

### EGC-2 and EGC-3 promote RNA interference

Given the roles of EGC-2 and EGC-3 in siRNA production, we then examined whether these proteins are required for RNAi response by feeding animals bacteria expressing dsRNAs targeting specific nematode genes. *pos-1* encodes a zinc-finger protein that is required for early embryonic cell fate decision[65]. RNAi targeting *pos-1* induces embryonic arrest in F1 embryos of animals exposed to dsRNA[65]. We fed the mutants with bacteria expressing dsRNAs targeting *pos-1* and found that *egc-3(-)* animals exhibited resistance to *pos-1*-mediated RNAi, whereas *egc-2(-)* animals responded similarly to wild-type animals (Figure. 7A). Animals lacking EGC-3 were also defective for experimental RNAi targeting *mex-3*, which encodes a KH domain protein that regulates the development of early *C. elegans* embryos (Figure. 7B)[66]. We further tested the silencing efficiency of a germline-expressed *gfp::h2b* transgene upon *gfp* RNAi. The loss of EGC-3 significantly prohibited the silencing effect of GFP::H2B upon feeding RNAi targeting *gfp* (Figure. 7C).

**Figure 7.**
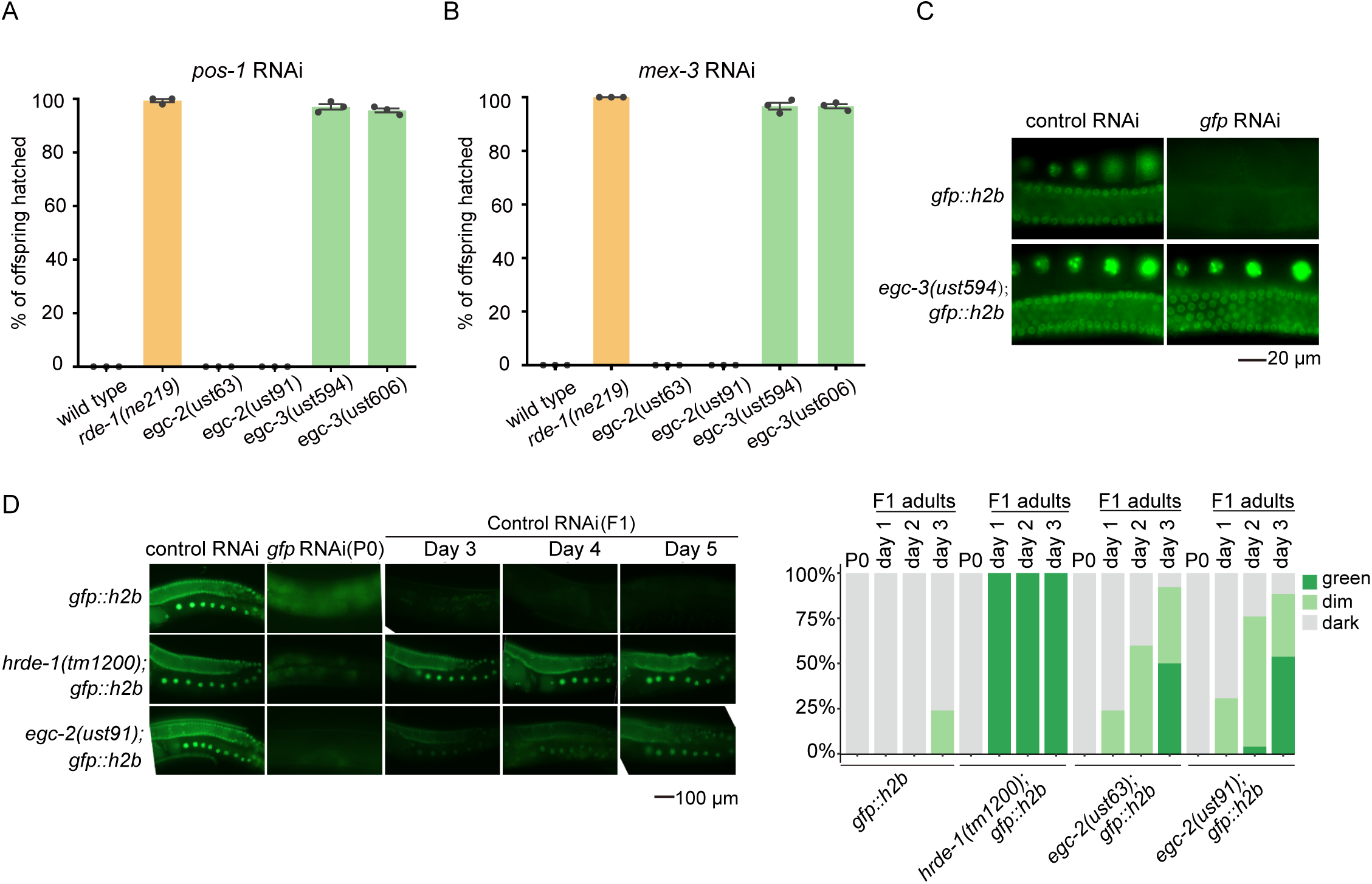
EGC-2 and EGC-3 promote RNA interference. (A and B) Bar graphs quantifying the number of hatched embryos following RNAi targeting *pos-1* and *mex-3* in the indicated animals. (C) Fluorescence images of the indicated animals subjected to or not subjected to *gfp* RNAi. EGC-3 is required for feeding RNAi targeting germline-expressed GFP. Animals expressing GFP::HIS-58 were exposed to *gfp* RNAi and bleached embryos were cultured on RNAi plates seeded with bacteria expressing *gfp* dsRNA. (D) Animals expressing GFP::HIS- 58 were exposed to *gfp* dsRNA and bleached F1 progeny were grown in the absence of dsRNA. GFP expression in P0 adult animals on day 1 and F1 adult animals on days 1, 2 and 3 was imaged by fluorescence microscopy with a 40 × objective. Adult day 1 was defined as the 24- h period after the hermaphrodite laid its first egg, and adult day 2 was defined as the subsequent 24-h period, etc [90]. The percentages of the indicated P0 and F1 animals expressing GFP were quantified. The data represent the scoring of at least 30 animals in each generation and for each genotype.

SID-1 and RDE-11 are required for dsRNA transportation and siRNA biogenesis during the RNAi response[67–69]. Recent studies reported that *meg-3/4* animals produce aberrant siRNAs targeting *sid-1* and *rde-11*, which silence the expression of these two genes and consequently result in defects in feeding RNAi response[45–47]. These two genes were also abnormally silenced in *egc-1* and *elli-1* mutants[26]. We found that siRNAs targeting these two genes are dramatically upregulated in *egc-3(-)* animals (Figure. S5A). We then performed mRNA-seq of wild-type and *egc-3(-)* animals. As expected, *sid-1* and *rde-11* mRNA levels were dramatically downregulated in *egc-3* mutants (Figure. S5B). Thus, these data suggested that the silencing of the *sid-1* and *rde-11* genes may underlie the defects of *egc-3(-)* animals in exogenous RNAi.

Interestingly, although the requirement for EGC-2 in experimental RNAi was less pronounced than that of EGC-3, the defect in RNAi response of *egc-2(-)* animals was more apparent during the inheritance phase of RNAi responses, suggesting that EGC-2 promotes the transgenerational inheritance of RNAi response (Figure. 7D). Together, these results suggest that EGC-2 and EGC-3 promote RNA interference in germ cells.

## Discussion

Here, via TurboID-based proximity labeling technology combined with RNAi-based genetic screening, we identify two novel intrinsically disordered proteins, EGC-2 and EGC- 3, that are required for E granule formation and small RNA homeostasis. The loss of EGC-2 or EGC-3 results in disruption of the perinuclear localization of the EGO complex and the PICS complex. Both EGC-2 and EGC-3 were enriched in the E granule. Small RNAomics revealed that EGC-2 and EGC-3 promote the production of 5’ E-class siRNA, but limit piRNA production. Moreover, EGC-3 is required for RNAi response and EGC-2 promotes RNA inheritance. Together, we speculate that EGC-2 and EGC-3 are key nodes in the core interaction network of E granule assembly, and are crucial for maintaining the homeostasis of the E granule and the enclosed small RNAs.

### E granule assembly

Biomolecular condensates are networks of protein‒protein interactions and protein‒RNA interactions, where high-valence scaffold proteins serve as central nodes that drive condensate formation. For example, G3BP proteins function as central nodes in stress granule (SG) assembly, exhibiting the highest centrality within the core SG network[70–72]. The interaction of G3BP with UBAP2L and Caprin-1 increases valency, facilitating liquid‒liquid phase separation (LLPS) and subsequent SG formation[73]. In this study, we identified two intrinsically disordered proteins, EGC-2 and EGC-3, as key players in E granule assembly. However, the molecular mechanisms underlying their roles in E granule assembly remain to be elucidated. Since the perinuclear localization of EGC-2 and EGC-3 is mutually dependent and the depletion of either protein disrupts the perinuclear localization of both the PICS and EGO complexes, we speculate that these two proteins may both function as central nodes or scaffold proteins during E granule assembly. Notably, EGC-2, but not EGC-3, has been identified as an interactor of TOFU-6 and EGO-1, suggesting that it may directly interact with the PICS and EGO complexes [26, 50, 51]. Based on these findings, we propose a model in which EGC-2 and EGC-3 may serve as central nodes in E granule assembly, with EGC-2 potentially interacting with the PICS and EGO complexes. EGC-3 may facilitate the recruitment of E-class RNA transcripts, thereby promoting the extension of the EGO module along RNA transcripts to generate siRNAs (Figure. 8). Future studies will be necessary to validate this hypothesis. Approaches such as immunoprecipitation‒mass spectrometry (IP‒MS) to identify EGC-2 or EGC-3 interactors could provide further insights into the molecular mechanisms of E granule assembly. Moreover, developing new techniques to profile RNA molecules within E granules may elucidate whether and how RNA transcripts contribute to the formation of E granules.

**Figure 8.**
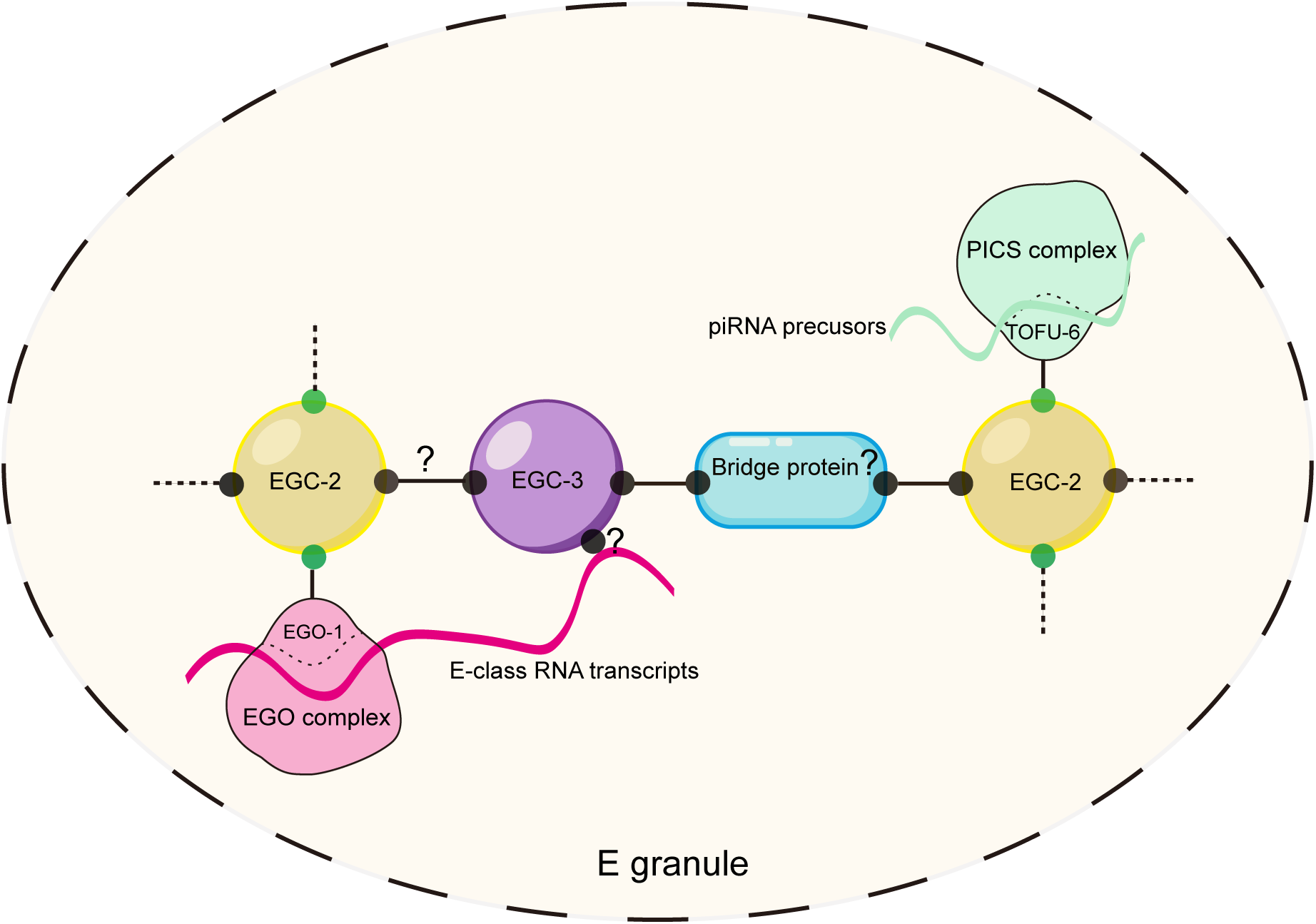
Working model. A model showing the roles of EGC-2 and EGC-3 in the core interaction network of E granule. EGC-2 and EGC-3 act as central nodes in the network. EGC- 2 interacts with the EGO and PICS complexes. EGC-3 may recruit E-class RNA transcripts to promote the elongation of the EGO module along the RNA transcripts to generate siRNAs. The potential interactions between EGC-2 and EGC-3, as well as the existence of bridging proteins that may mediate their interaction, need to be further investigated.

### The distinct functions of E granule assembly factors in the production/function of different classes of small RNAs

Biomolecular condensates are microscale compartments in eukaryotic cells that lack surrounding membranes, but enclose biomolecules including proteins and nucleic acids. Yet, whether the positioning of biomolecules in these membraneless orgalleles is essential for corresponding biological processes is still ambugious. For example, P body formation is considered a consequence, rather than a driver, of miRNA-mediated gene silencing[74]. Manipulating condensation without disrupting enzymatic activity or RNA binding can disentangle effects due to the disruption of condensates versus the loss of RNP complex activity[75]. In this study, through candidate-based RNAi screening, we identified two intrinsically disordered proteins, EGC-2 and EGC-3, that act as upstream regulators of E granule assembly. The biogenesis of 5*’* E-class siRNAs was significantly impaired in *egc-2* or *egc-3* mutants, although this loss was less pronounced in *egc-2* mutants than in those lacking the EGO module or EGC-3. These results suggest that either E granules still form in *egc-2(-)* animals, albeit to a degree not detectable via light microscopy, or that EGC-3 directly contributes to E-class siRNA production, in addition to promoting E granule assembly. Further investigations into the sequence characteristics and processing patterns of RNA intermediates in E granules as well as the proteins binding to these transcripts, will provide deeper insights into the role of E granules in siRNA amplification.

Intracellular condensates are highly multicomponent systems that enclose various RNP bodies, undertaking diverse RNA processing events or sequential biological reactions[1]. For instance, in the germline cells of the *Drosophila* egg chamber, many proteins involved in piRNA processing are enriched in a unique perinuclear structure, the nuage, including the following: Vasa, Aub, AGO3, Krimper, Maelstrom, Spindel-E, Tejas, Vreteno, and so on[76]. These proteins function sequentially to generate primary piRNAs or ping-pong cycle-derived secondary piRNAs[77]. Interestingly, the proteins involved in the ping-pong cycle exhibit a hierarchical genetic interaction for their localization to the nuage, which could be an indication of their sequential order of function in piRNA biogenesis[78–80]. Similarly, in *C. elegans* germ cells, many piRNA biogenesis factors are enriched in germ granules, including the E granule-enriched PICS/PETISCO complex, Z granule-enriched PARN-1 and P granule-enriched HENN-1[31, 50, 51, 81, 82]. Additionally, the *C. elegans* PIWI protein PRG-1 is present in both the Z and P granules, with a preference for the Z granule[31]. Our findings indicate that E granule assembly defects generally do not diminish mature piRNA abundance, suggesting that enrichment of the PICS complex in E granules is not essential for piRNA processing. Intriguingly, disrupting E granule assembly or E granule localization of the EGO complex results in a slight increase in piRNA abundance and an overactive piRNA silencing system. Although the underlying molecular mechanisms are not fully understood, the data imply that the enclosing of both the PICS and EGO complexes to E granules might provide a strategy for regulating piRNA abundance or the piRNA silencing system. A recent study reported that *parn- 1* mutants accumulate 3*’* untrimmed piRNA precursors, which promotes the generation of EGO- 1-dependent anti-piRNAs, that regulate the function of PRG-1 in 22G RNA production[83]. Thus, the positioning of piRNA processing factors to distinct germ granule compartments may provide a means of monitoring particular steps of piRNA processing. Further studies are needed to examine whether and how the perinuclear localization of other piRNA processing factors within germ granules regulates piRNA production and the piRNA surveillance pathway.

## Materials and Methods

### C. elegans strains

The Bristol strain N2 was used as an untagged control in proximity labeling assay and the standard wild-type strain in RNA-seq analysis. All other strains used in this study were generated by genome editing or genetic crosses and are listed in Table S2. Unless otherwise indicated, the animals were cultured at 20°C according to standard methods[84].

### Construction of transgenic and mutant strains

For the TurboID transgene, the coding sequence of TurboID fused to a 6AA (amino acid) linker sequence (GGAGGTGGAGGTGGAGCT) was inserted upstream of the *elli-1* stop codon via the CRISPR/Cas9 genome editing method[31, 55]. TurboID sequence was PCR amplified from the plasmid pAS31[54]. Left and right homologous arms were PCR amplified from N2 genomic DNA. The vector backbone was PCR amplified from the plasmid pCFJ151. All these fragments were joined together by Gibson assembly to form the repair plasmid using the ClonExpress MultiS One Step Cloning Kit (C113-02, Vazyme). The injection mixture contained the repair plasmid (50 ng/μL), pDD162 (50 ng/μL), co-injection marker pSG259 (5 ng/μL) and four sgRNAs targeting 3*’ elli-1* sequence (30 ng/μL of each sgRNA). Three days later, F1 animals expressing pharyngeal GFP were isolated. After another three days, the targeted animals with TurboID insertion were screened via PCR.

For the *egc-2::gfp::3xflag* ectopic transgene, the DNA elements were fused and integrated into *C. elegans* chromosome II (ttTi5605 locus) via the MosSCI method[85]. For *in situ* tagging of *egc-3*, the coding sequence of *gfp::3xflag* was inserted upstream of the *egc-3* stop codon via the CRISPR/Cas9 method. Plasmids containing repair templates were generated using the ClonExpress MultiS One Step Cloning Kit. The method of obtaining integrated animals through microinjection and PCR screening was the same as above.

For deletion mutants, three or four sgRNAs (30 ng/μL of each sgRNA) were co-injected into N2 animals with pDD162 (50 ng/μL), pSG259 (5 ng/μL). Three days later, F1 animals expressing pharyngeal GFP were isolated. After another three days, the deletion mutants were screened via PCR as previously described[56].

### Fluorescent streptavidin staining

Synchronized L1 animals were first cultured at 15℃ to late L4 stage, and then at 25°C until young adult, since culturing animals at higher temperatures (25°C) increases biotinylation activity[86, 87]. Young adult animals were suspended and washed three times with M9 buffer to remove excess bacteria. The animals were then transferred to CXY buffer (0.4 × M9, 0.1 M NaN3) and dissected on poly-L-Lysine-Prep Slides (E678002, Sangon Biotech). The dissected animals on the slides were treated with 3.3% (w/v) paraformaldehyde fixation solution in PBS at room temperature for 15 min, followed by two washes with PBS. Slides were then fixed for 15 min in – 20°C methanol and washed twice in PBST, after which 1:1000 streptavidin-Alexa Fluor 488 (S11223, Invitrogen) in PBST was added to the slides. The slides were kept in a humidified chamber overnight at 4°C. The slides were then washed four times with PBST and two times with PBS. Each wash lasted 5 minutes. Anti-fade Mounting Medium (E675011, Sangon Biotech) was added to the slides and covered with a coverslip. The slides were sealed with nail polish and kept at 4°C until imaging.

### Streptavidin-HRP blotting

Whole worm lysates were prepared by boiling animals at 70°C for 10 min in 4x LDS sample buffer (P0731, Beyotime), and then separated on 4–12% Bis-Tris gels (LK307, Epizyme), transferred onto NC membranes, and probed with HRP-conjugated Streptavidin 1:4000 (D111054, Sangon Biotech) for detection via highly sensitive ECL luminescence reagent (C500044, Sangon Biotech).

### Streptavidin affinity pull-down and mass spectrometry analysis

Approximately 70000 synchronized L1 animals were placed on NGM plates seeded with concentrated OP50 food and cultured as described previously[34]. Young adults were collected and washed with M9 buffer more than three times to remove excess biotin. Animals were then resuspended in RIPA buffer (50 mM Tris-HCl (pH 8.0), 150 mM NaCl, 1% Triton X-100, 0.1% SDS, and 1 mM EDTA in ddH2O) supplemented with cOmplete protease inhibitor cocktail (Roche) and solution P. The solution P contained 5 mg/10 mL pepstatin A and 0.1 M PMSF. The resuspended pellets of the animals were then subjected to liquid nitrogen freezing and grinding cycles until they were ground into powder. The lysates were subsequently centrifuged at 13200× RPM for 30 minutes. Zeba Spin Desalting Columns (89891, Thermo) were used for desalination and removal of biotin. The supernatant was mixed with Streptavidin magnetic beads (88816, Thermo) at a ratio of 40 µl beads/500 µl proteins and incubated overnight at 4°C with constant rotation. Beads were then washed for 5 min, two times with RIPA buffer, once with 1 M KCl, once with 0.1 M Na2CO3, and once with 2 M urea in 10 mM Tris-HCl (pH 8.0). Beads were subsequently resuspended in PBS and subjected to on-beads trypsin digestion. The bound proteins on the beads were dissolved in 50 mM Tris-HCl (pH 8.0) supplemented with 8 M urea and 5 mM DTT and incubated at 37°C for 1 hour with shaking. After incubation, 1 M iodoacetamide (Sigma) was added to a final concentration of 15 mM, and the samples were incubated for an additional 30 minutes in the dark at room temperature. The samples were then diluted with three volumes of 50 mM Tris-HCl (pH 8.0) to allow for trypsin digestion. 1ug sequencing grade-modified trypsin (V5111, Promega) was mixed with the samples and incubated overnight at 37°C with shaking. The reaction was quenched by adding 2% FA (Sigma) for acidification. The supernatant was taken out and concentrated for LC‒MS analysis.

### Candidate-based RNAi screening

All RNAi assays were performed at 20°C by placing synchronized embryos on RNAi plates as previously described[88]. HT115 bacteria expressing the empty vector L4440 (a gift from A. Fire) were used as controls. Bacterial clones expressing double-stranded RNAs (dsRNAs) were obtained from the Ahringer RNAi library and sequenced to verify their identity. All feeding RNAi assays were performed for two generations except for sterile animals, which were RNAi-treated for one generation.

### RNA isolation and sequencing

Synchronized young adult animals were incubated with TRIzol (Invitrogen) for five quick liquid nitrogen freeze‒thaw cycles before isopropanol precipitation. For 21U-RNA sequencing, RNA samples were subjected to DNase I digestion (Thermo Fisher) and re-extracted via the TRIzol method. For 22G-RNA sequencing, samples were further treated with RNA 5′- polyphosphatase (Epicentre) as previously described[26].

The prepared RNA samples were subjected to deep sequencing via an Illumina platform (Novogene Bioinformatic Technology Co., Ltd.). Briefly, small RNAs ranging from 17 to 30 nt were gel purified and ligated to a 3′ adaptor (5′-pUCGUAUGCCGUCUUCUGCUUGidT-3′) and a 5′ adaptor (5′-GUUCAGAGUUCUACAGUCCGACGAUC-3′). The ligation products were gel purified, reverse transcribed, and amplified via Illumina’s sRNA primer set (5′- CAAGCAGAAGACGGCATACGA-3′; 5′-AATGATACGGCGACCACCGA-3′). The samples were then sequenced via the Illumina HiSeq platform.

### RNA-seq analysis

For 22G-RNA analysis, clean reads ranging from 17 to 30 nt, processed by Novogene, were mapped to the *C. elegans* transcriptome assembly WS243 as previously described[26]. For mature 21U-RNA analysis, clean reads were mapped to the mature piRNA regions and the transcriptome assembly WS243 respectively[50]. The numbers of reads targeting each transcript were counted using custom Perl scripts. The normalization value was generated by subtracting the number of total reads corresponding to sense rRNA transcripts from the number of total reads mapped to the transcriptome. A cutoff criterion of a twofold change was applied to identify the differentially expressed small RNAs. Scatter plots and Venn diagrams were generated via custom R or Python scripts and modified in Adobe Illustrator. 22G RNA reads were aligned to the *C. elegans* genome WBcel235 via Bowtie2 v.2.2.5 with default parameters, and IGV v.2.5.3 was used to visualize the alignment results.

### Microscope and images

To image young adults, the animals were immobilized in ddH2O with 0.5 M NaN3 and mounted on 2% agarose pads. To image germ cells and embryos, the animals were dissected in CXY buffer (0.4×M9, 0.1 M NaN3) on a coverslip and then mounted on 1% agarose pads. Images were collected using a Leica upright DM4 B microscope equipped with a Leica DFC7000 T camera. For stitched images, individual images were collected by shifting the slides horizontally and then stitched manually in overlapping regions.

### Statistics

The bar graphs with error bars represent the means +/- SDs. All experiments were performed with independent animal samples or the indicated number of replicates. Statistical analysis was performed with a two-tailed Student’s t test or an unpaired Wilcoxon test as indicated. Origin 2023 or R scripts were used for statistical analysis.

### Data Availability

The raw sequence data reported in this paper have been deposited in the Genome Sequence Archive in the National Genomics Data Center (China National Center for Bioinformation/Beijing Institute of Genomics, Chinese Academy of Sciences) under accession codes CRA019966, CRA019967 and CRA019969.

## Supporting information

Table S1.putative E granule components identified by TurboID proximity labeling

Table S2.C.elegans strains used in this study

## Acknowledgments

We are grateful to the members of the Guang laboratory for their comments. We are grateful to Dr. Gang Wan’s laboratory for the suggestions and technique support. We are grateful to the International *C. elegans* Gene Knockout Consortium and the National Bioresource Project for providing the strains. Some strains were provided by the CGC, which is funded by the NIH Office of Research Infrastructure Programs (P40 OD010440).

## Funding

This work was supported by grants from the National Key R&D Program of China (2022YFA1302700) and the National Natural Science Foundation of China (32230016, 32270583, 32470633, 32400435, 2023M733425 and 32300438), the Research Funds of Center for Advanced Interdisciplinary Science and Biomedicine of IHM (QYPY20230021) and the Fundamental Research Funds for the Central Universities.

## Author contributions

K.L., S.G., and X.C. conceptualized the research; K.L., X.F., S.G. and X.C. designed the research; K.L., K.W., and X.F. performed the research; K.L., K.W., X.F., Xiaona.H., C.Z., and Xinya.H. contributed new reagents/analytic tools; K.L., X.F., Q.W., S.G. and X.C. wrote the paper.

## Competing interests

The authors declare no competing interests.

## Supplementary Information

Figures S1 to S5 Tables S1, S2

## Supplemental figure legends

**Figure S1.**
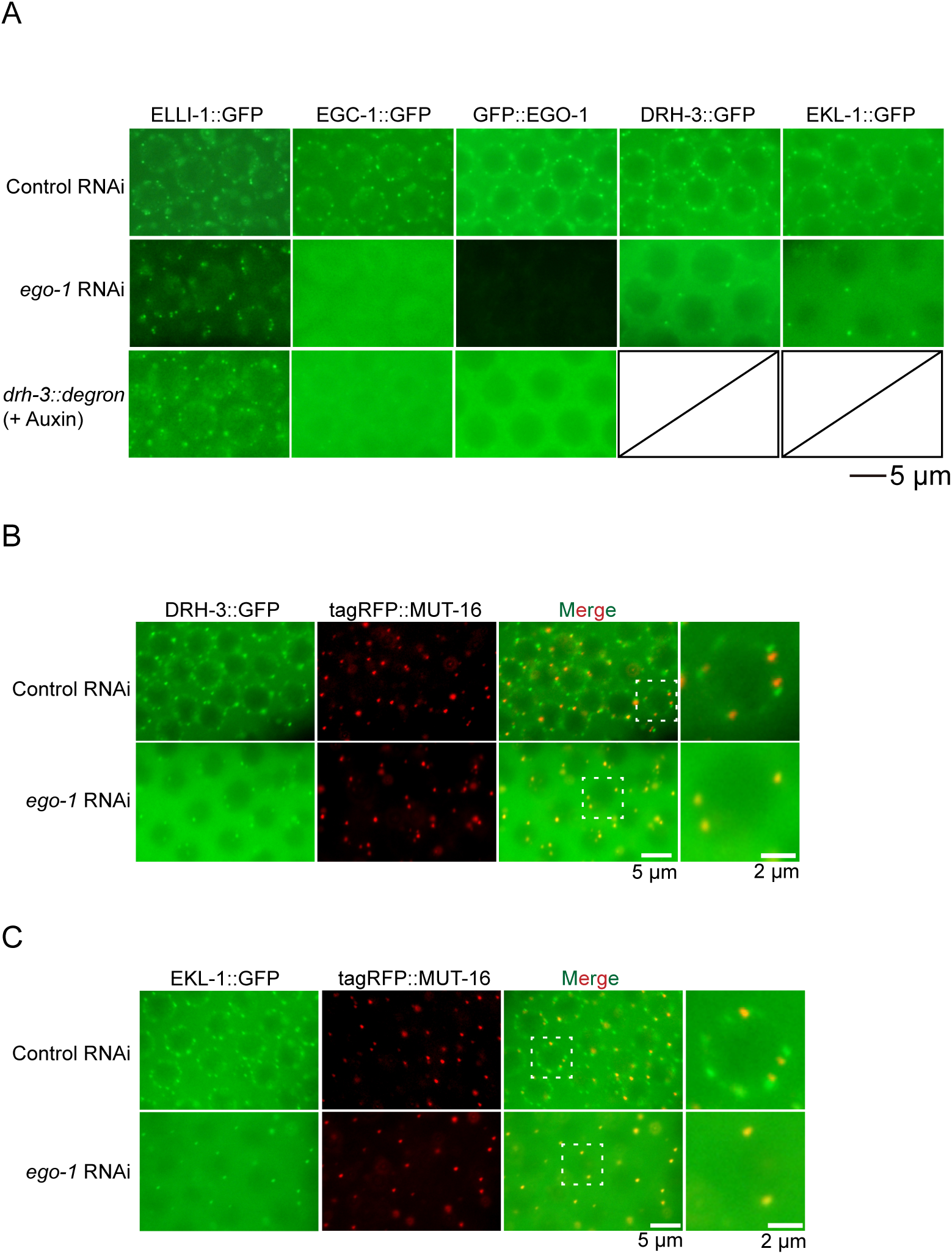
ELLI-1 is required for the E granule localization of other EGO complex components. (A) Fluorescence micrographs of the GFP-tagged EGO complex components in adult germ cells from the indicated animals. RNAi knockdown of *ego-1* disrupted the localization of EGC-1 and EGO-module factors but did not disrupt ELLI-1 foci. Depletion of *drh-3* via the auxin-inducible degron (AID) system disrupted EGC-1 and EGO-1 foci but did not disrupt ELLI-1 foci [1]. (B and C) Fluorescence micrographs of pachytene germ cells expressing DRH-3::GFP and tagRFP::MUT-16, or EKL-1::GFP and tagRFP::MUT-16 following feeding RNAi targeting *ego-1*. All images are representative of more than three animals.

**Figure S2.**
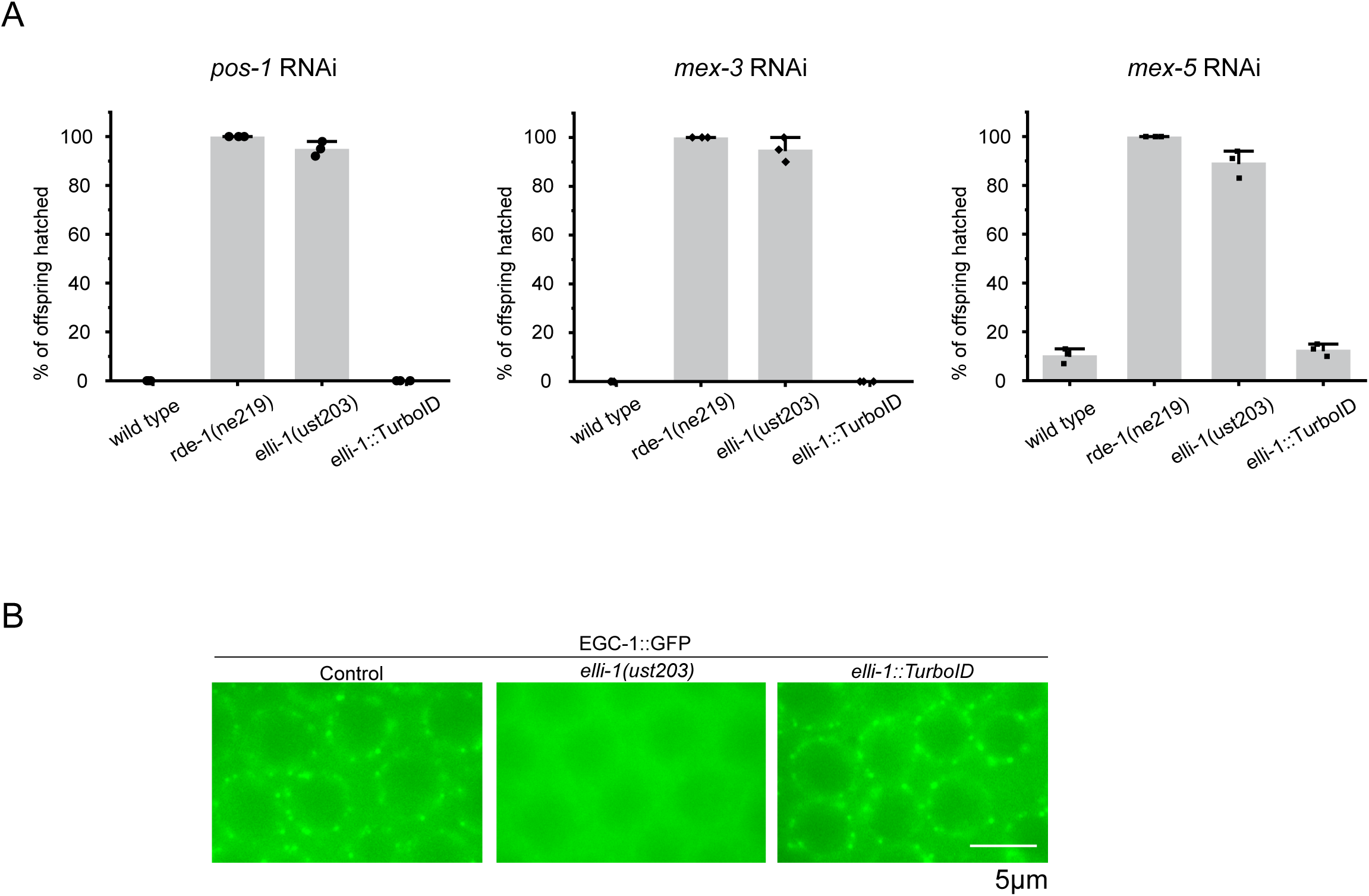
*elli-1::TurboID* produces functional fusion proteins. (A) Bar graphs showing the quantification of hatched embryos after feeding RNAi targeting *pos-1*, *mex-3* and *mex-5* in the indicated animals. Synchronized embryos were grown on P0 RNAi plates with bacteria expressing *pos-1*, *mex-3* and *mex-5* dsRNA. Three L4 P0 animals were picked to F1 plates and laid eggs. Total F1 embryos and hatched F1 embryos were scored. The *rde-1(-)* and *elli-1(-)* animals were defective for *pos-1*, *mex-3* and *mex-5* RNAi. Tagging ELLI-1 with TurboID did not affect ELLI-1 function. Data are shown as mean±SD of three biological replicates. (B) Fluorescence micrographs showing pachytene germ cells of wild-type, *elli-1(ust203)*, and *elli-1*::TurboID animals expressing EGC-1::GFP. Tagging ELLI-1 with TurboID did not disturb the perinuclear localization of EGC-1::GFP.

**Figure S3.**
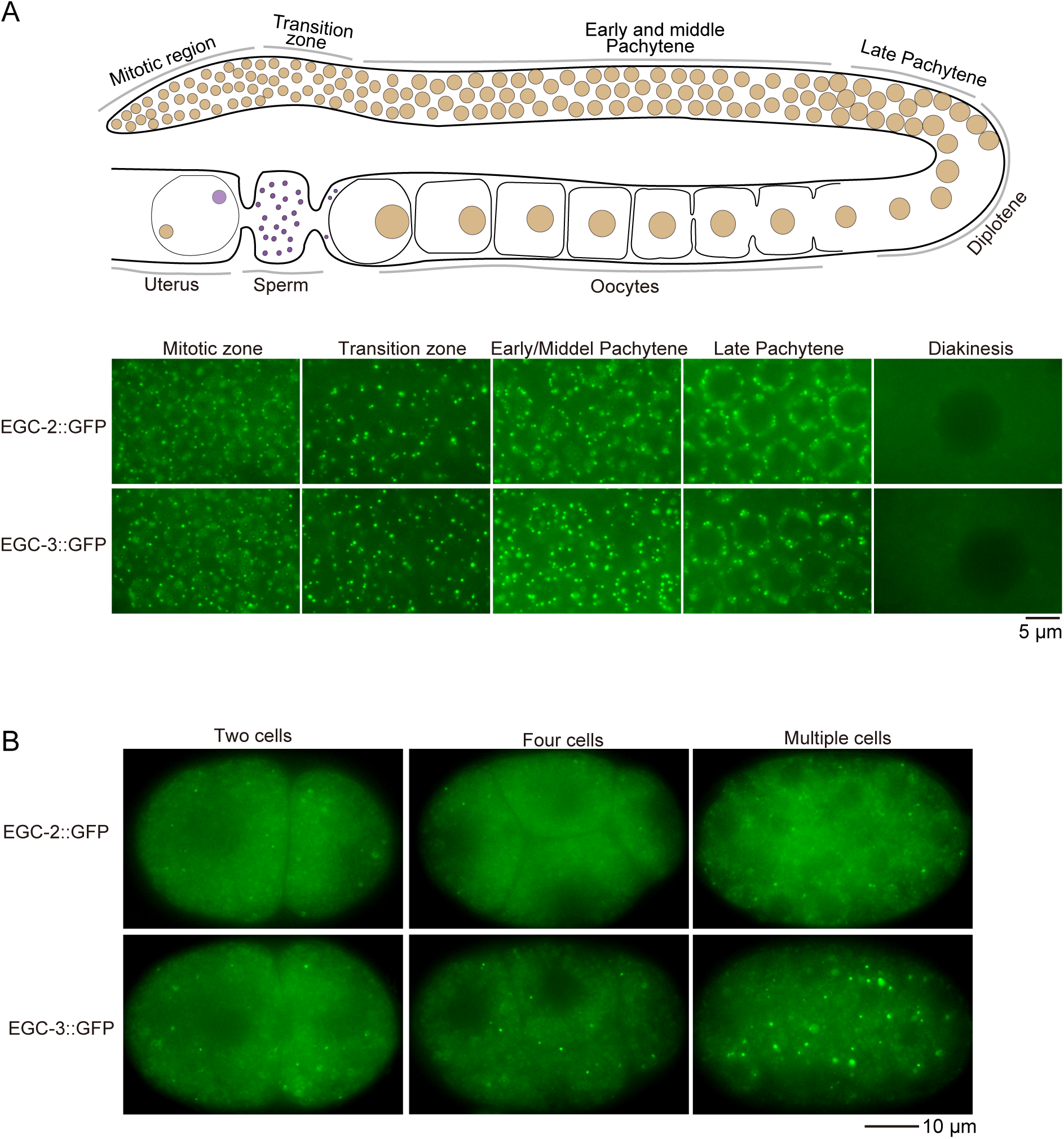
The expression patterns of EGC-2 and EGC-3. (A) Fluorescence micrographs of EGC-2::GFP and EGC-3::GFP in adult germ cells at different differentiation stages. Upper, a schematic of the *C. elegans* adult hermaphrodite germline. Bottom, fluorescence micrographs of the germ cells at different stages. (B) Fluorescence micrographs of early embryos expressing EGC-2::GFP and EGC-3::GFP.

**Figure S4.**
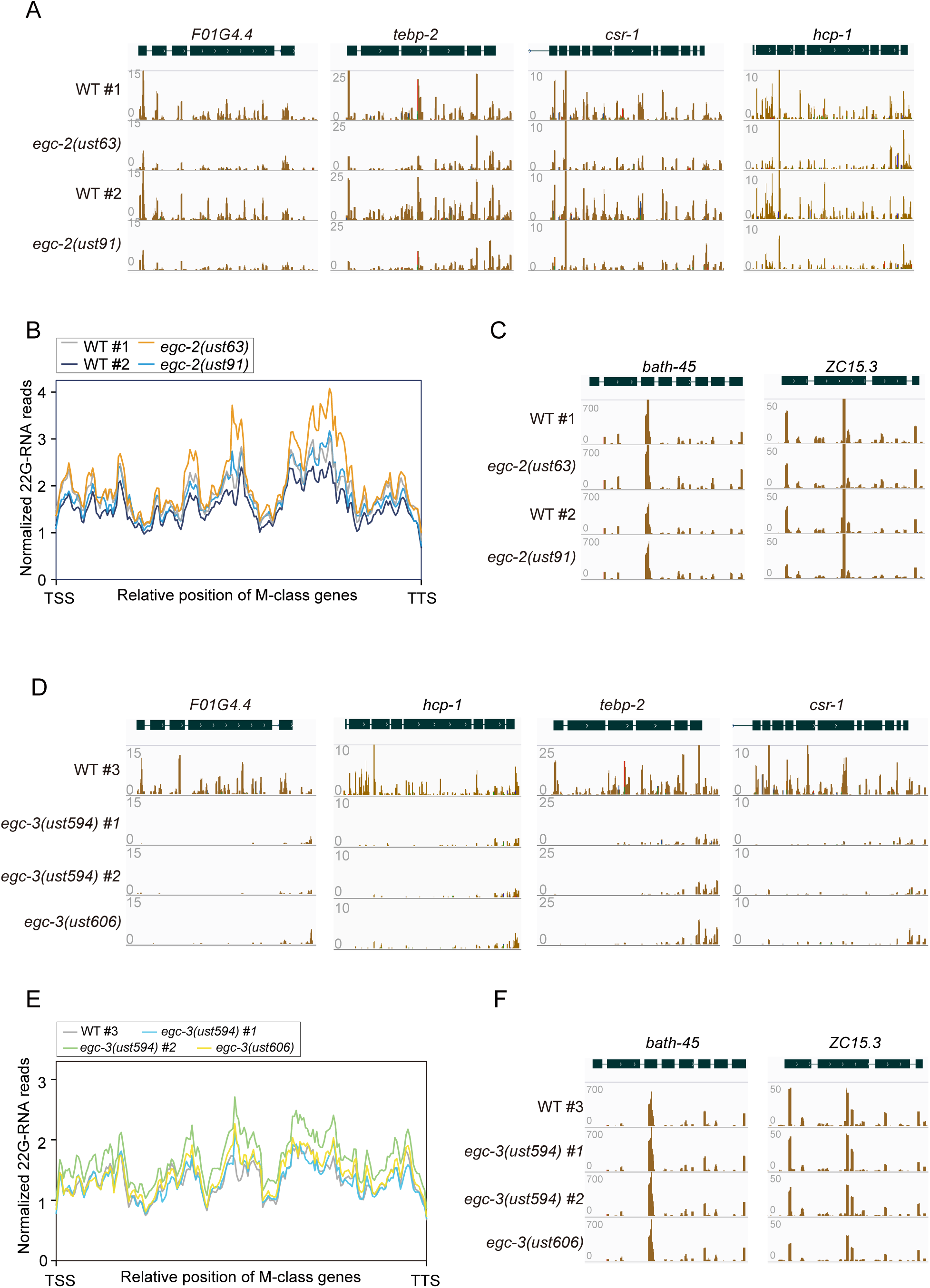
EGC-2 and EGC-3 promote specialized synthesis of 5’ E-class siRNAs. (A and D) Normalized 22G RNA read distribution across E-class siRNA target genes in the indicated animals, including *F01G4.4*, *tebp-2*, *csr-1* and *hcp-1*. (B and E) Metaprofile analysis showing the distribution of normalized 22G RNA reads (RPMs) along M-class genes in the indicated animals. (C and F) Normalized 22G RNA read distribution across M-class siRNA target genes in the indicated animals, including *bath-45* and *ZC15.3*. Loss of EGC-2 or EGC-3 did not affect the production of M-class 22G RNAs.

**Figure S5.**
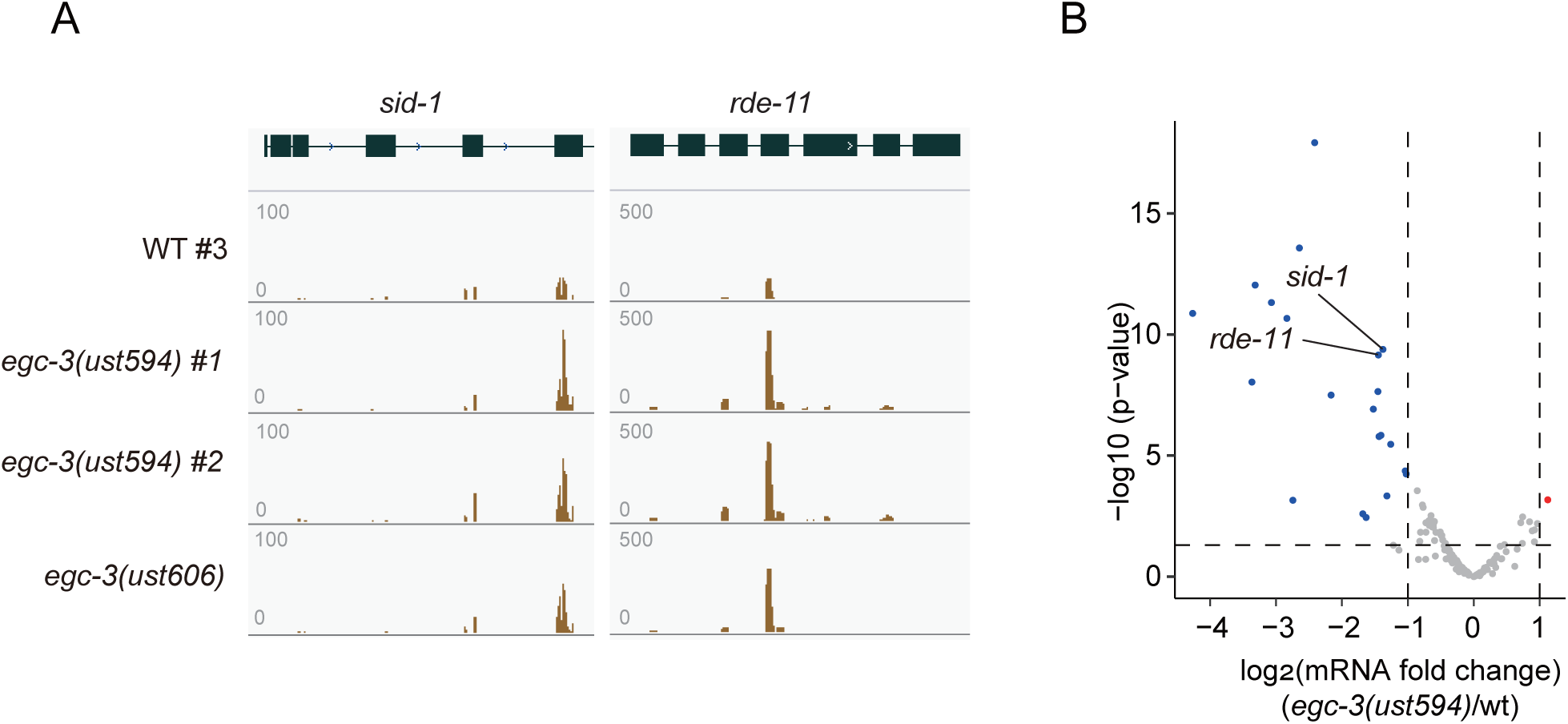
EGC-3 regulates the expression levels of the RNAi-related genes *sid-1* and *rde-**11***. (A) Normalized 22G RNA read distribution across *sid-1* and *rde-11* in the indicated animals. 22G siRNAs mapping to *sid-1* and *rde-11* were both upregulated in *egc-3(-)* animals. (B) Volcano plot showing the fold change of mRNAs of 199 genes with upregulated siRNAs in *egc-1 or elli-1* animals[2]. *sid-1* and *rde-11* are indicated.

## Notes

### Competing Interest Statement

The authors have declared no competing interest.

